# NPC1 confers metabolic flexibility in Triple Negative Breast Cancer

**DOI:** 10.1101/2022.05.05.490674

**Authors:** KI O’Neill, LW Kuo, MM Williams, HE Lind, LS Crump, NG Hammond, NS Spoelstra, MC Caino, JK Richer

## Abstract

Triple negative breast cancer (TNBC) often undergoes at least partial epithelial-to-mesenchymal transition (EMT) to facilitate metastasis. Identifying EMT-associated characteristics can reveal novel dependencies that may serve as therapeutic vulnerabilities in this aggressive breast cancer subtype. We find that *NPC1*, which encodes the lysosomal cholesterol transporter Niemann-Pick Type C1 is highly expressed in TNBC as compared to estrogen receptor-positive (ER+) breast cancer and is significantly elevated in high grade disease. We demonstrate that *NPC1* is directly targeted by microRNA-200c (miR-200c) a potent suppressor of EMT, providing a mechanism for its differential expression in breast cancer subtypes. Silencing of *NPC1* in TNBC causes an accumulation of cholesterol-filled lysosomes and drives decreased growth on soft agar and invasive capacity. Conversely, overexpression of NPC1 in an ER+ cell line increases invasion and growth on soft agar. We further identify TNBC cell lines as cholesterol auxotrophs, however, they do not solely depend on NPC1 for adequate cholesterol supply. Genetic inhibition of *NPC1* in TNBC cell lines led to altered mitochondrial function and morphology, suppression of mTOR signaling, and accumulation of autophagosomes. A small-molecule inhibitor of NPC1, U18666A, decreased TNBC proliferation and synergized with the chemotherapeutic drug, paclitaxel. This work suggests that NPC1 promotes aggressive characteristics in TNBC and identifies NPC1 as a potential therapeutic target.

## INTRODUCTION

Triple negative breast cancer (TNBC) is an aggressive subtype representing 15% of newly diagnosed breast cancer cases. Because TNBC often acquires resistance to chemotherapy (30-50% of patients)^(1)^, identification of targetable vulnerabilities is imperative. Compared to estrogen-receptor positive (ER+) and human epidermal growth factor amplified (HER2+) breast cancers, TNBC have a rapid peak rate of recurrence as metastatic disease, often within 3-5 years of initial diagnosis^(2)^. TNBC have significantly lower expression of the microRNA miR-200 family than ER+ BC and normal breast epithelium^(3,4)^. This miRNA family, known as the “guardian of the epithelial phenotype”, potently suppresses epithelial-to-mesenchymal transition (EMT)^(3,5-7)^. Silencing or deletion of miR-200c by methylation or micro-deletions allows EMT to occur by permitting translation of developmental, mesenchymal, and neuronal genes not typically expressed in normal epithelial cells^(8-11)^. Work with human TNBC cell lines and mouse mammary carcinoma models demonstrated that restoration of miR-200c to TNBC drives a switch to a more epithelial, less invasive, and less metastatic phenotype, through direct targeting of mesenchymal transcription factors including ZEB1, a suppressor of E-cadherin and other epithelial markers. Restoration of miR-200c to TNBC can thus can reveal potential dependencies of TNBC.

In addition to repressing genes traditionally associated with EMT, restoration of miR-200c to TNBC cells reduces levels of immune-modulatory and metabolic genes^(12,13)^. We discovered that *NPC1*, encoding the lysosomal cholesterol transporter Niemann-Pick Type C-1, was among the genes most significantly reduced by miR-200c, along with additional changes to genes involved in cholesterol and lipid metabolism^(12)^, suggesting cholesterol transport as a potential pathway of interest in TNBC progression. Cellular cholesterol is tightly regulated and can be derived from *de novo* biosynthesis or lysosome-mediated uptake. Following endocytosis of cholesterol-containing low density lipoproteins (LDL) from the microenvironment, cells process cholesterol within the lysosome and utilize NPC1 to efflux cholesterol to the endoplasmic reticulum^(14)^. NPC1 is best studied in the context of loss-of-function mutations that lead to the Niemann-Pick Type C lysosomal storage disorder, characterized by abnormal accumulation of cholesterol within the lysosome, driving trafficking defects and lysosomal dysfunction^(15-17)^. NPC1 is also the mechanism by which mTOR senses cholesterol levels and serves as a component of the lysosomal mTOR signaling scaffold^(18,19)^.

In the context of cancer, NPC1 activity and function is understudied. Here, we report that NPC1 may serve as a potential therapeutic target in TNBC, where it is elevated compared to ER+ BC. We find that restoration of miR-200c directly targets *NPC1* and represses NPC1 protein levels. Suppression of NPC1 in TNBC leads to slowed proliferation, decreased anchorage-independent survival and decreased invasion, and altered mitochondrial function. Thus, NPC1 may serve as a potential therapeutic target in TNBC.

## METHODS

### Cell culture and reagents

MDA-MB-231 and BT549 cell lines were purchased directly from ATCC. Sum159PT (RRID:CVCL_5423) cells were purchased from the UCCC Tissue Culture Core, and the MCF7 cells were obtained from Dr. Kate Horowitz at University of Colorado Anschutz Medical Campus. Cell lines were authenticated by Short Tandem Repeat DNA Profiling (Promega) and tested for mycoplasma in the University of Colorado Cancer Center (UCCC) Cell Technologies shared resource (September 2020). For culture, MDA-MB-231 (MDA-231) cells were grown in MEM with 5% FBS, 1mM HEPES, 2mM L-glutamine, and insulin; BT549 (NCI-DTP Cat# BT-549, RRID:CVCL_1092) cells were cultured in RPMI 1640 medium with 10% FBS, and insulin; MCF7 In MEM with 5% FBS and insulin; Sum159PT cells were grown in Ham F-12 with 5% FBS, penicillin/streptomycin, hydrocortisone, insulin, HEPES, and L-glutamine supplementation. Prior to experimentation, all cells were switched to DMEM +10% FBS, which was used throughout all experiments unless otherwise noted.

#### Stable NPC1 knockdown and exogenous expression cell lines

pLVX-NPC1(WT)-FLAG (RRID:Addgene_164972) was a gift from R. Zoncu at University of California, Berkeley. pcDNA3.1 (RRID:Addgene_79663) and pcDNA3.1-NPC1 were from Genscript. pLKO-empty and pLKO encoding NPC1-targeted shRNAs were purchased from The University of Colorado-Anschutz Functional Genomics Core, with shRNA #1 targeted toTRCN0000005428 (3’UTR) and shRNA #2 targeted to TRCN0000418552 (CDS). Cells were selected in neomycin (500μg/mL) or puromycin (1μg/mL) for ∼2 weeks prior to experimentation. Notably, we struggled to maintain the BT549 shNPC1 cell lines long-term, and BT549 and MDA-MB-231 shNPC1 cells were not consistently viable after freeze-thaw.

### Transfections

#### miR-200c experiments

Negative, scrambled control (4464058, Thermofisher Scientific) or miR-200c (4464066,MC11714,Thermofisher Scientific) mimics were used at a final concentration of 50 nmol/L. All transfections were done using either RNAi Max or Lipofectamine 3000 (Thermo Fisher Scientific) and experiments were conducted following the manufacturer’s protocols.

#### Luciferase assay

50nM of scramble mimic negative control or miR-200c-3p mimic was co-transfected with 1μg of Pmir-glo (E1330,Promega) fused with NPC1-3’UTR containing seed regions of miR-200c binding site or mutant. Cells were lysed after 48hrs of transfection. The luciferase activities were measured by the Dual-Luciferase® Reporter Assay System (E1910, Promega) according to the manufacture instruction with internal Renilla control. Site-direct mutagenesis of seed region was performed utilizing GeneArt™ Site-Directed Mutagenesis PLUS System (A14604,Invitrogen). Primers used are listed in Supplementary Table 1. N*PC1 siRNA silencing:* Transient siRNA transfections were done using 30-60μM NPC1 siRNA from Ambion/Thermofisher (catalog AM16704, ID 106016), in combination with Lipofectamine RNAi Max (ThermoFisher) and conducted according to manufacturer’s protocols.

### qRT-PCR

Total RNA was isolated using TRIzol RNA Isolation (Qiagen) and cDNA was synthesized with the qScript cDNA SuperMix (QuantaBio). qRT-PCR was performed in an ABI 7600 FAST Thermal Cycler. SYBR Green quantitative gene expression analysis was performed using ABsolute Blue qRT-PCR SYBR Green Low ROX Mix (Thermo Fisher Scientific). Primers are listed in Supplementary Table 2.

### Western Immunoblotting

Whole-cell protein extracts (20-40 μg) were denatured, separated on SDS-PAGE gels, and transferred to polyvinylidene fluoride membranes. After blocking in 5% non-fat milk in Tris-buffered saline–Tween, membranes were probed overnight at 4°C. The following antibodies are used in this study: NPC1 (Novus Biologicals, NB400); LDLR (Novus Biologicals NBP1-06709); HMGCR (ThermoFisher PA5-37367); HMGCS1 (Thermo Fisher Scientific Cat# PA5-29488, RRID:AB_2546964); DHCR24 (Cell Signaling Technology Cat# 2033, RRID:AB_2091448); LAMP1 (Abcam Cat# ab25630, RRID:AB_470708); Phospho-S6 Kinase (Thr389) (Cell Signaling, #9205); SQSTM1/P62 (Cell Signaling #5114); LC3B (Cell Signaling, #3868). Secondary antibodies were Licor IR Dye Goat anti-rabbit 800 IgG (#926-32211) or goat anti-mouse 680 (#926-68070).

### Nutrient Deprivation

Lipoprotein-depleted fetal bovine serum (LPDS, Kalen Biomedical #880100) and used at 5% or 10% in DMEM, and when noted supplemented with 10μg/mL Human LDL (Kalen Biomedical 770200-4). For MBC:Cholesterol complexing, methyl-β-cyclodextrin (Medchem Express, HY-101461) and cholesterol (sigma C8667) were warmed to 67°c, combined at 1:1 molar ratio (1% MBC to 20μg/mL Chol), added to 37°c media, and vortexed. For amino acid starvation, EBSS was supplemented with 5mM glucose, 1mM sodium pyruvate, and 2mM glutamine.

### Dil-LDL Uptake

Cells seeded 24-48hr in 24 well dishes were washed 1x with PBS and starved in 5% LPDS in DMEM for 1 hour, then given 10μg/mL dil-LDL (Kalen Biomedical # 770230-9) in 5% LPDS/DMEM and incubated for ∼5 hours at 37°c. Following lysis in RIPA, cells were spun for 5min at 20,000g. 5μL supernatant was transferred to 96-well plate reader (Biotek) and read at 520/580nm. Remaining supernatant was used to measure protein by BCA assay, and uptake was normalized to μg of protein.

### 2D and 3D growth and Invasion assays

*For U18666A IC-50* measurements, BT549, MDA-MB-231, and Sum159PT cells were seeded in 96 well dishes at 6k, 5k and 3k cells per well, respectively. Cells were given serial dilutions of U18666A for 48 hours and compared to 1% DMSO control. Cells were fixed in 10% neutral buffered formalin for 20 minutes, washed with PBS, and incubated in crystal violet stain for 20 minutes before rinsing with diH_2_O. Plates were dried overnight and stain was resuspended in 5% acetic acid, absorbance was measured at 570nm.

*For U18666A and Paclitaxel combination*, BT549 cells were seeded in 96well dishes at 6k cells/well. The next day, 3 doses of paclitaxel (2, 7, 12nM) and 3 doses of U18666A (2, 4, 8μM) were given alone or in combination with n=3 wells each. After 48 hours, crystal violet was performed. Additive/synergy calculations were done using SynergyFinder 2.0 (SynergyFinder, RRID:SCR_019318).

*Confluency over time* of NPC1 siRNA cells was conducted using xCELLigence RTCA (Agilent Technologies). Cells were seeded in xCELLigence plates 24 hours following transfection and “cell index” was measured every 60 minutes for 96 hours.

*For 3D/soft agar assays*, cells were plated in triplicates in a 6-well plates in 0.5% bottom agar and 0.3% top agar containing 5,000 cells/well (Sum159PT) or 30,000 cells/well (MCF7). Cells were grown for approximately 14 days with once-weekly media change and stained with nitro blue tetrazolium at harvest. For LPDS soft agar experiments, agar contained 5% LPDS-DMEM, with or without 10μg/mL LDL and these concentrations were maintained in top media. For MBC:Cholesterol experiments, 5% FBS-DMEM was given vehicle or 20μg/mL cholesterol in 1:1 molar ratio complex with MBC, as described previously. Experiments were performed three separate times, imaged using Optronix GelCount, and quantified using ImageJ software (ImageJ, RRID:SCR_003070).

#### Invasion Assay

3D transwell invasion assays were performed with 8.0µm pore transwell inserts in 24 well plates. Filters were coated overnight with 200µg/mL Cultrex UltiMatrix Basement Membrane Extract (R&D Systems, Cat. # BME001). 50,000 cells per well were then seeded in serum free media. Invasive cells were assessed 24 hours later via fixation with 10% NBF and staining with 0.1% crystal violet.

### Microscopy

#### Mitochondrial Immunocytochemistry and quantification

Cells were grown on glass coverslips for 48 hours, then fixed in 10% formalin (*Fisher Scientific Cat# SF93*), permeabilized with 0.1% Triton-X100, and blocked in 10% normal goat serum with 0.3M glycine in PBS. Fixed cells were incubated overnight with mouse anti-mitochondria (MTC02, Thermo Fisher Scientific Cat# MA5-12017, RRID:AB_10983622) diluted at 1:500, washed in PBS 3 times, followed by 45 min at room temperature with anti-mouse Alexa Fluor 568 secondary antibody (Thermo Fisher Scientific Cat# A-11004, RRID:AB_2534072) diluted 1:500 and slides were mounted using Prolong Diamond Antifade Mountant DAPI (Thermo Fisher P36966). For qualitative scoring of cells according to mitochondrial morphology, imaging was done using an Olympus BX40 microscope with 100x/ objective lens and FITC filter.

Random fields (25-30) were analyzed and each cell was classified into 3 categories according to the main mitochondrial morphology (small, medium, elongated). For automated analysis of mitochondrial size, slides were scanned in a Zeiss LSM780 confocal microscope with 63X oil objective. Z-stack from 5 random fields were acquired with a slice thickness of 0.45 microns, and the built-in 3D function on the Zeiss software was used to render 3D images of the mitochondrial network surface at 1,4, and 8X zoom. Images were analyzed in ImageJ (RRID:SCR_003070) to obtain the average mitochondria size per cell, from ∼20 cells per condition.

#### Filipin Stain

Cells were grown on glass coverslips and fixed in 10% formalin, quenched with 100mM Glycine. 0.05mg/ml Filipin (Santa Cruz, #480-49-9) was used in 10% normalized goat serum in PBS and stained for 1.5hr, mounted with Prolong Diamond Antifade Mountant, and imaged using Olympus BX40 microscope and 40x objective.

#### NPC1 IHC

Formalin fixed paraffin embedded tissues were cut at 5μm, baked, and deparaffinized in a series of xylenes and graded ethanols. Antigens were heat retrieved using citrate buffer, and tissues were stained for NPC1 using Novus Bio #NB400-148 primary antibody followed by goat anti-rabbit HRP, and DAB (Vector Labs, RRID:AB_2819346).

### Seahorse Assay

The Seahorse Mitochondrial stress test was performed according to manufacturer’s instructions using Seahorse XFe96 format (Agilent). The morning of the experiment, cells, media was changed to Seahorse XF DMEM medium, pH 7.4 (103575-100) with 10mM glucose, 2mM glutamine, and 1mM sodium pyruvate. Media was also refreshed immediately prior to assay run. Basal oxygen consumption rate (OCR) was measured prior to addition of mitochondrial inhibitors: Oligomycin 2μM, carbonilcyanide p-triflouromethoxyphenylhydrazone (FCCP) 1.5μM, and Rotenone + Antimycin A 2μM each. Spare respiratory capacity was measured as the change between OCR before and after FCCP addition. All measurements were normalized to cell count, obtained through crystal violet of seahorse plate immediately following Seahorse assay.

### Mitochondrial Reactive Oxygen Species

Mitochondrial ROS was determined in plate-reader format using mitoSOX red (ThermoFisher M36008). Cells were stained in Hank’s Balanced Salt Solution without phenol red, with 5μM mitoSOX for 25 minutes. For positive control wells, 10μM Antimycin A was added 10 minutes prior to removal of mitoSOX staining. Cells were washed 1x PBS and read at 510/580 using Biotek plate reader, then fixed for crystal violet staining. Fluorescent intensity was normalized to unstained control wells and cell count via crystal violet.

## RESULTS

### NPC1 is significantly elevated in TNBC and is directly targeted by miR-200c

Query of *NPC1* in publicly available patient datasets demonstrated higher *NPC1* mRNA in patients with basal, ER-, and high-grade tumors relative to ER+ low grade disease (**Figure 1A, Supplementary 1A**). By western blot, TNBC cell lines had higher expression of NPC1 relative to ER+ breast cancer cell lines (**Figure 1B, Supplementary 1B**), and immunohistochemistry demonstrated NPC1 expression in both primary tumor and lung metastasis in an MMTV-driven mouse model of breast cancer (**Supplementary 1C**). The regulation of NPC1 in cancer has not been fully elucidated, but in normal physiology, NPC1 is induced through multiple transcription factors, including cholesterol-mediated Sterol Response Element Binding Proteins (SREBPs)^(20,21)^, and the cAMP response element binding protein (CREB) in steroidogenic cells^(22)^. However, expression of *NPC1* in breast cancer data sets does not correlate with genes in these pathways (**Supplementary 1D**), suggesting that NPC1 is not upregulated due to broadly increased cholesterol or lysosomal signaling.

**Figure 1:**
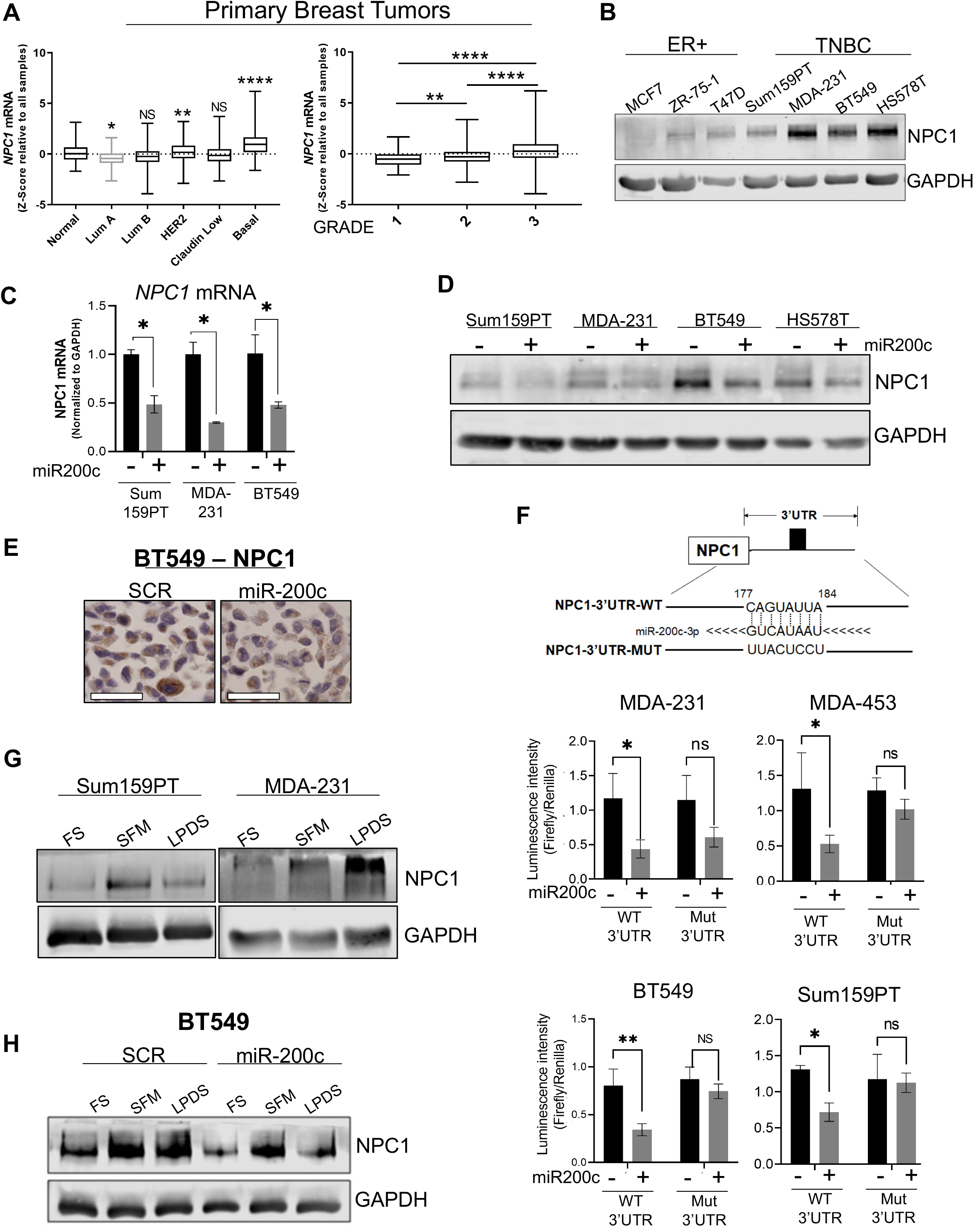
*NPC1* is significantly elevated in aggressive breast cancers and is directly regulated by miR-200c. **A)** *NPC1* in 2019 METABRIC breast cancer cohort by subtype (left, ordinary one-way ANOVA relative to “normal”) and grade (right, ordinary one-way ANOVA). **B)** Western blot of baseline NPC1, panel of breast cancer cell lines. **C)** Effect of miR-200c on *NPC1* in 3 TNBC cell lines by qPCR, after 48 hours transfection. Unpaired T-Test **D)** Effect of miR-200c on NPC1 following 72 hours transfection. **E)** Immunohistochemistry of NPC1 in BT549 -/+ miR-200c, scale bar=50µm. **F)** Luciferase assay performed on wildtype (WT) or mutated (Mut) *NPC1* 3’UTR, with and without exogenous miR-200c. (Student’s T-Test) **G)** Western blot of NPC1 in serum free media (SFM) or lipoprotein depleted serum (LPDS) for 24 hours, in two TNBC cell lines. **H)** Effect of miR-200c on NPC1 induction by 24hr SFM and LPDS treatment. miR-200c transfection=96 hours. P-values denoted by asterisks where * ≤ 0.05; ** ≤ 0.01; *** ≤ 0.001; ****≤ 0.0001.

Functionally confirming our prior gene array data^(12)^, exogenous restoration of miR-200c suppressed NPC1 at the mRNA and protein level in multiple TNBC cell lines (**Figure 1C-1E**). The 3’UTR of *NPC1* contains a predicted binding site for miR-200c (targetscan.org)^(23)^; therefore we tested for evidence of direct binding and regulatory activity at this site. Luciferase reporter assays confirmed direct binding of miR-200c to the wildtype but not mutated *NPC1* 3’UTR predicted binding sequence (**Figure 1F**). Notably, miR-200c had no effect on LDLR mRNA or protein levels (**Supplementary 1E-F**). These data reveal a novel miR-200c/NPC1 regulatory axis in breast cancer that may contribute to the differences in patient tumors observed in publicly available datasets.

As in normal glandular epithelial cells^(21)^, we observe that TNBC cells increase NPC1 in response to low cholesterol (lipoprotein-depleted serum, LPDS) and serum-free media (SFM) conditions (**Figure 1G**). Although miR-200c attenuates total levels of NPC1 at baseline, it was unable to completely suppress NPC1 induction in low-serum and low-sterol conditions (**Figure 1H, Supplementary 1G**), demonstrating that miR-200c collaborates with other metabolic factors to regulate NPC1 levels. Together, these data show increased expression of NPC1 in TNBC, influenced by the loss of inhibitory miR-200c, and suggest basal NPC1 upregulation as a unique feature in these tumor cells.

### NPC1 supports breast cancer cell invasion and growth in soft agar

Based on the expression of NPC1 in breast cancer and its known roles in relevant cholesterol metabolism and mTOR signaling pathways, we sought to determine if NPC1 supports tumor-promoting characteristics *in vitro*. Following shRNA knockdown of NPC1 using two unique shRNAs (**Figure 2A**), TNBC cells demonstrate cholesterol accumulation within the lysosomes, as visualized with filipin stain (**Figure Supplementary 2A**). These cells grow at a slowed rate in 2D culture compared to control cells (**Figure 2B, Supplementary 2B**). In soft agar, which recapitulates anchorage-independent growth, NPC1 silencing decreased the number and size of colonies in Sum159PT cell lines (**Figure 2C**). TNBC cell lines were less invasive when NPC1 was silenced (**Figure 2E**). This is consistent with a recent study demonstrating that NPC1 inhibition reduces migration and invasion in Chinese hamster ovarian and A431 squamous carcinoma cell lines^(24)^.

**Figure 2:**
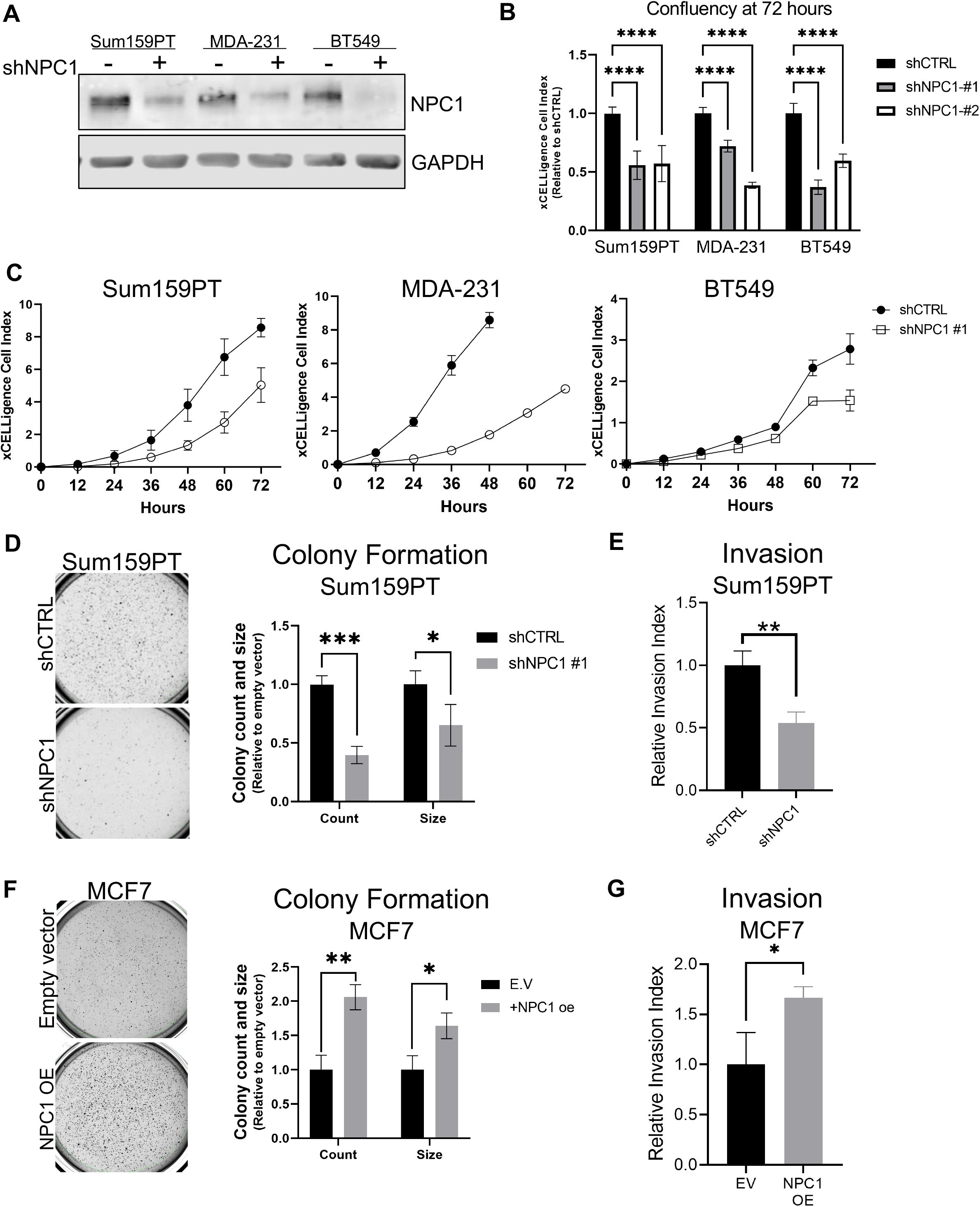
NPC1 supports breast cancer cell invasion and growth in soft agar. **A)** Western blot confirmation of NPC1 knockdown by shRNA. **B)** Relative confluency of 3 TNBC cell lines with two unique shRNAs, following 72 hours in 2D cultured, as measured by xCELLigence. **C)** Growth of cell lines (“shRNA#1”) over 72 hours, as measured by xCELLigence. **D)** Sum159PT colony formation in soft agar, two weeks after seeding. **E)** Invasion of TNBC cells through Cultrex after 24hrs, from serum free DMEM toward 10% DMEM. **F)** Colony formation in soft agar over 14 days and **G)** invasion through Cultrex gel over 24hrs. MCF7 cells transfected with empty vector (pcDNA3.1) or constitutively active NPC1 (pcDNA3.1-NPC1). All statistics: students T-Test.

Strikingly, exogenous expression of NPC1 in ER+ MCF7 cells led to increased number and size of colonies in soft agar (**Figure 2F, Supplementary 2E**), and increased invasion of +NPC1 cells after 24 hours (**Figure 2G**). The effect of NPC1 on anchorage-independent growth and invasiveness demonstrates regulation of multiple tumor cell-intrinsic properties relevant to the metastatic potential of TNBC.

### TNBC cells are cholesterol auxotrophs but do not solely depend on NPC1 for adequate cholesterol supply

In the absence of adequate LDL-derived cholesterol from the microenvironment, most cell types in normal physiology are able to produce cholesterol *de novo* through the cholesterol biosynthesis pathway^(14,25,26)^. However, recent studies identified “cholesterol auxotrophy”, the inability to survive without exogenous cholesterol, in varying types of cancer^(27,28)^,suggesting cholesterol uptake as a targetable pathway. NPC1 specifically transports lysosomal cholesterol to the endoplasmic reticulum^(29)^ where cellular cholesterol levels are sensed and controlled ^(17,30)^. By depleting the endoplasmic reticulum of cholesterol, NPC1 inhibition affects both supply of exogenous cholesterol as well and disrupts cholesterol homeostasis^(29,31)^. How loss of NPC1 alters cholesterol homeostasis in BC has not been evaluated. Further, the dependency of breast cancer on exogenous cholesterol has not been reported.

To determine if disruption of cholesterol supply and homeostasis plays a role in the slowed growth observed in Figure 2, cholesterol auxotrophy in breast cancer was evaluated. A panel of breast cancer cell lines was grown in LPDS with or without supplemental LDL for 7 days, leading to significantly slowed growth in 7 of 13 cell lines (**Figure 3A**). This occurred disproportionately in TNBC (5/6) as compared to non-TNBC (2/7). These trends were maintained during 14 day culture (**Supplemental 3A, top**). Of note, cholesterol auxotrophs grew normally for ∼3-4 days, and then experienced a loss of viability over subsequent days (**Supplemental 3A, bottom**). Because metabolic demands can differ in 2D versus 3D conditions, cholesterol dependence was tested over 14 days in soft agar, which demonstrated that cell lines maintain cholesterol dependence/independence in 3D anchorage-independent culture (**Figure 3B**).

**Figure 3:**
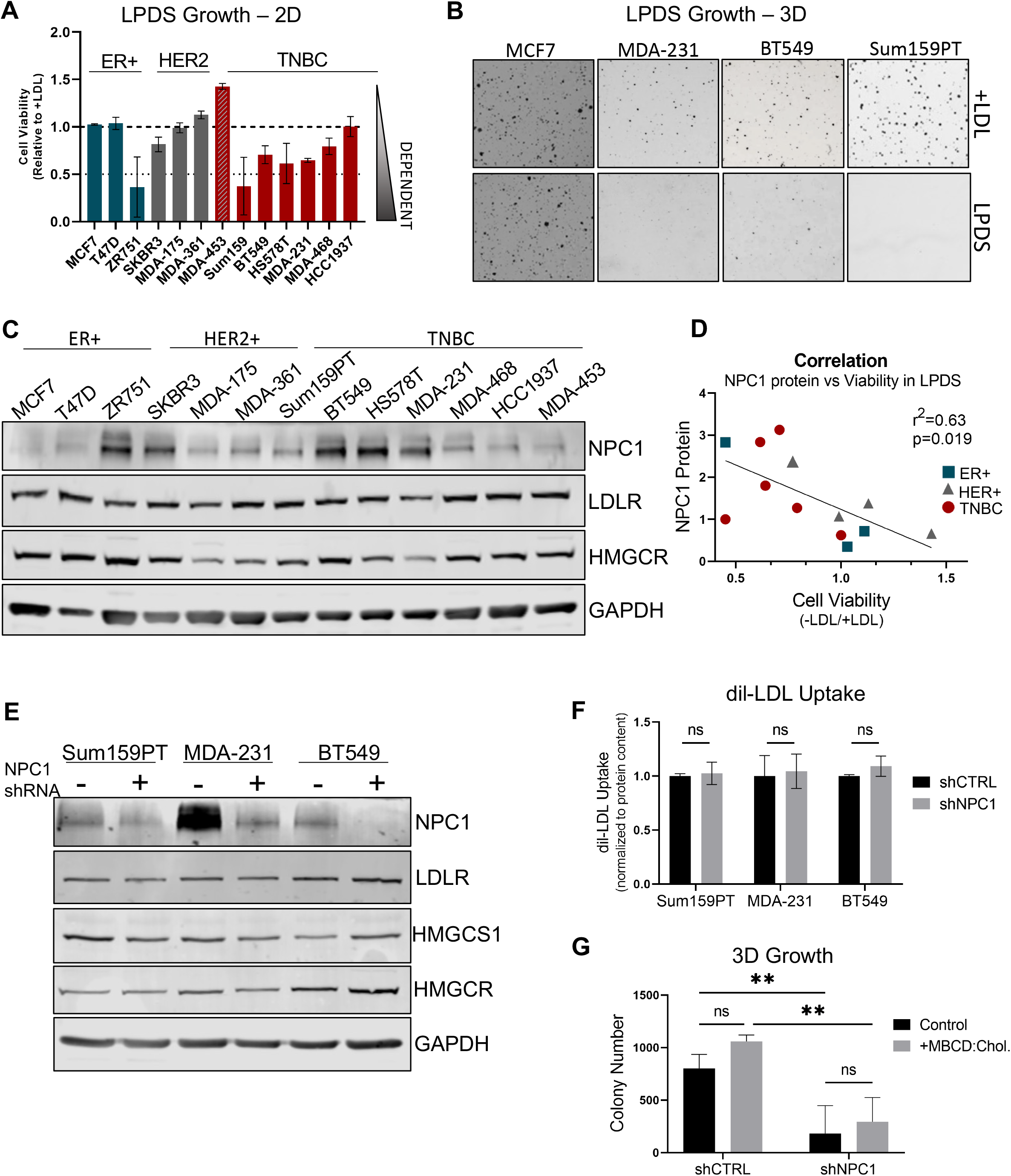
TNBC cells are cholesterol auxotrophs but do not solely depend on NPC1 for adequate cholesterol supply. **A)** Growth of breast cancer cell lines over 7 days in cholesterol-deplete (5% LPDS) relative to cholesterol-replete (5% LPDS + 10µg/mL LDL) media. **B)** Colony formation in soft agar, cholesterol-deplete (*top*) and –replete (*bottom*, 10µg/mL LDL);14 days. **C)** Basal levels of key cholesterol metabolism proteins NPC1, LDLR, and HMGCR in a panel of breast cancer cells. **D)** Correlation of NPC1 protein levels (quantified from western, 3C) to cell viability in LPDS (viability score from 3A). **E)** Western blot of key cholesterol proteins in shRNA#1 cell lines. **F)** Uptake of fluorescent dil-LDL following 6 hour incubation, analyzed by plate reader and normalized to protein content. **G)** Growth of Sum159PT cells in soft agar when supplemented with MBCD:Cholesterol (1% MβCD complexed with 10µg/mL cholesterol) over 14 days.

In other models of cholesterol auxotrophy, which include ALK+ lymphomas and renal cell carcinomas, the phenotype was driven by low expression or mutations in key cholesterol biosynthesis genes^(27)^. However, loss-of-function mutations within these genes were not identified in breast cancer cell lines (**Supplemental 3B**), and cholesterol auxotrophy does not correlate with mRNA levels of any cholesterol biosynthesis or uptake-related genes in these cell lines (**Supplemental 3C**). While cholesterol auxotrophy weakly correlates (Pearson r= -0.63, p=0.019) with NPC1 protein expression (**Figure 3C, 3D**), the most strongly correlated genes had limited relation to cholesterol metabolism (**Supplementary 3D-3F**). These data suggest that while cholesterol auxotrophy exists in a subset of breast cancer, it may not be driven by a single, global mechanism. However, protein expression of NPC1 is positively correlated with dependency on exogenous cholesterol and may play a role in maintaining adequate cholesterol supply in cells that cannot produce it *de novo*.

To evaluate if NPC1 affects and supports cholesterol metabolism, key cholesterol readouts were evaluated following NPC1 knockdown. Low cholesterol conditions drive activation SREBP-1 and -2, which transcriptionally upregulate a large set of sterol-responsive enzymes. These include the Low-Density Lipoprotein Receptor (*LDLR*) and the rate-limiting step of cholesterol biosynthesis, HMG-CoA reductase (*HMGCR*)^(32)^. While cholesterol-related genes were differentially expressed in control versus shNPC1 cells, changes did not trend in one direction nor were they consistent between cell lines (**Supplemental 4A-4C**). At the protein level, neither LDLR nor HMGCR were altered (**Figure 3E**) and the rate of LDL uptake was unchanged (**Figure 3F**), demonstrating that NPC1 loss does not specifically activate low-sterol responses in TNBC.

It remained possible that loss of lysosome-ER cholesterol transport could result in a detrimental level of cholesterol depletion in NPC1 knockdown cells, given abnormal patterns of cholesterol localization in NPC disease^(33)^. To address this, soft agar growth media was supplemented with cholesterol complexed to Methyl-β-Cyclodextrin (MBCD), which delivers cholesterol independently of the lysosome^(29,34)^. However, MBCD:cholesterol failed to rescue growth of NPC1 knockdown cell lines (**Figure 3G**).

Together, these data demonstrate the novel finding of cholesterol auxotrophy in TNBC cancer cells, which correlates with NPC1 protein expression. However, NPC1 silencing did not cause dramatic changes to TNBC cholesterol metabolism, nor did supplementation of cell-permeable cholesterol rescue cell growth. Thus, loss of NPC1-dependent cholesterol trafficking is likely not the mediator of the observed phenotype following NPC1 silencing in TNBC.

### NPC1 supports mitochondrial respiration and maintains mitochondrial morphology

In models of Niemann-Pick disease, fibroblasts and stem cells demonstrate abnormal mitochondrial metabolism^(16,35,36)^. However, the mechanisms driving mitochondrial defects in this disease remain poorly understood. In TNBC cells, NPC1 silencing suppressed basal respiration and spare respiratory capacity (**Figure 4A, Supplementary 5A-C**), while overexpression in ER+ MCF7 cells supported increased basal respiration (**Figure 4B**). Basal glycolysis was not affected by NPC1, but knockdown cell lines were less able to upregulate glycolysis following treatment by oligomycin, which inhibits mitochondrial ATP synthase (complex V). This suggests a limited ability to upregulate glycolysis to compensate for mitochondrial inhibition (**Supplementary 5D-5E**). Fitting with loss of mitochondrial respiration, NPC1 knockdown cells had increased basal levels of mitochondrial reactive oxygen species (ROS) (**Figure 4D**) and a greater relative increase in mitochondrial ROS when treated with Antimycin A, which is known to produce mitochondrial ROS due to electron transport chain complex III inhibition^(37)^ (**Supplementary 6A**).

**Figure 4:**
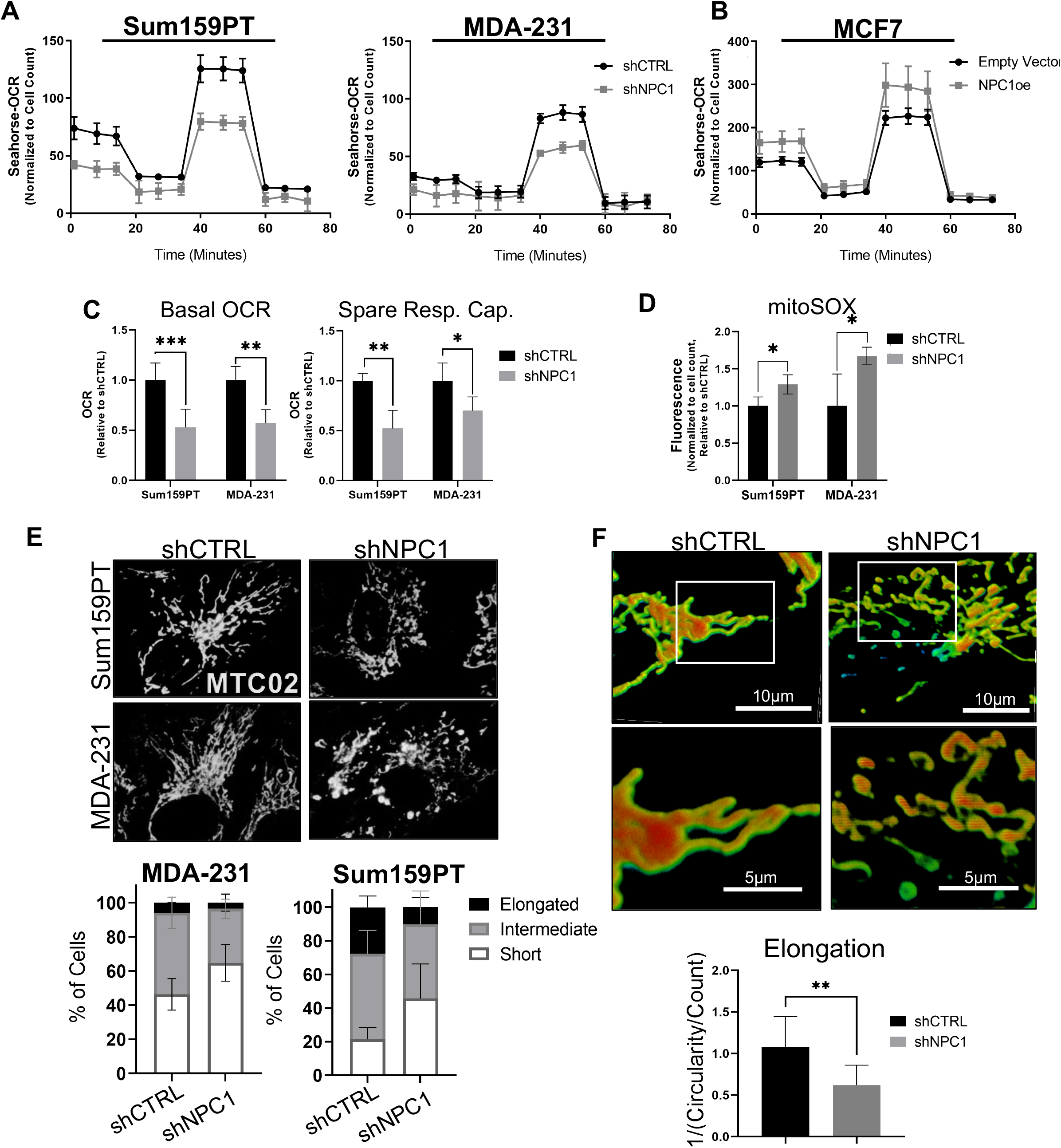
NPC1 supports mitochondrial respiration and maintains mitochondrial morphology. **A)** Oxygen Consumption Rate (OCR) during Seahorse mitochondrial stress test, performed on Sum159PT and MDA-231 cell lines with NPC1 knockdown. **B)** Mitochondrial stress test performed on MCF7 with pcDNA3.1-NPC1 or empty vector. **C)** Quantification of basal OCR and Spare Respiratory Capacity of TNBC cells from mitochondrial stress test shown in A,. **D)** Mitochondrial ROS as evaluated using mitoSox staining. **E)** *Top:* Representative images of fluorescence microscopy of TNBC cells stained with MTC02 (mitochondrial antibody). *Bottom:* quantification of cells with mitochondrial morphology defined as either elongated, intermediate, or short. **F)** Confocal Z-stacking of mitochondria stained with MTC02, with representative images of “elongated” (shCTRL) versus “shortened” (shNPC1) mitochondria. 60x with 4x zoom (top panel) and 60x with 8x zoom (bottom panel). Bottom: quantification of mitochondrial elongation, as quantified by ImageJ.

NPC1 loss-of-function has also been associated with altered mitochondrial morphology^(35)^, which could play a role in the observed decrease in oxygen consumption rate, cell growth, and invasion of NPC1 knockdown cells. Indeed, NPC1 silencing in TNBC cells increased the percentage of cells exhibiting “short” and “intermediate” mitochondria while reducing the percentage of “elongated” mitochondria (**Figure 4E and 4F, Supplementary 6B, 6C**). Mitochondrial morphology can be regulated by fission and fusion processes, however, no baseline changes to key fission/fusion proteins were detected. Interestingly, however, NPC1 expression altered fission/fusion signaling (MFN-1 and MFF) in response to metabolic and autophagic stress (**Supplementary 7A**). Mitochondria characterized as “shortened” in shNPC1 cell lines included “donut” mitochondria (**Figure 4F**), a morphology that is associated with decreased ATP production, defective calcium signaling, and increased mitochondrial ROS^(38,39)^.

Together, these data demonstrate that NPC1 silencing impairs mitochondrial ATP production as evaluated by oxygen consumption rate, which is associated with shortened mitochondrial morphology and increased mitochondrial reactive oxygen species (ROS). Given the key role of mitochondrial function in cell growth and metastasis^(40)^, defective mitochondria may contribute to the suppression of invasion and growth of TNBC cells following NPC1 inhibition.

### NPC1 affects mTOR and autophagy signaling under stress conditions in TNBC

NPC1 acts as the cholesterol “sensor” for mammalian target of rapamycin (mTOR), the central regulator of metabolic-related proliferative signaling. In normal physiology and under cholesterol-replete conditions, NPC1 recruits mTOR to the lysosomal surface, where mTOR signaling is initiated^(19)^. In the absence of cholesterol, altered conformation of NPC1 prevents this recruitment, prohibiting mTOR signaling^(18)^. In normal fibroblasts, neurons, and stem cells, silencing of NPC1 allows constitutive activation of mTOR even under cholesterol-depleted conditions. However, mTOR signaling in normal nutrient conditions is not affected by NPC1 silencing in these cell types^(18,35)^. mTOR/AKT is activated in numerous cancers, generally through mutations in upstream regulators including PTEN and PIK3CA^(41)^. Given the role of mTOR in cancer and its physical interaction with the lysosome and NPC1^(18,19,35)^, western blot was used to evaluate how NPC1 silencing affects mTOR signaling.

In Sum159PT cells, loss of NPC1 caused a reduction in the mTOR target pS6k (**Figure 5A**). This suggests a regulatory role for NPC1 in the context of TNBC, unique from that in other published models^(18,35)^. To establish if NPC1 still serves as a cholesterol “sensor” for mTOR in cancer models, control versus shNPC1 cells were starved of cholesterol using methyl-beta-cyclodextrin (**Figure 5A**). As has been published in HEK293T cells^(35)^, cholesterol starvation suppressed pS6K in control cells but not shNPC1 cells, demonstrating that in these contexts, mTOR is sensitive to low cholesterol and depends on NPC1 to sense cholesterol levels.

**Figure 5:**
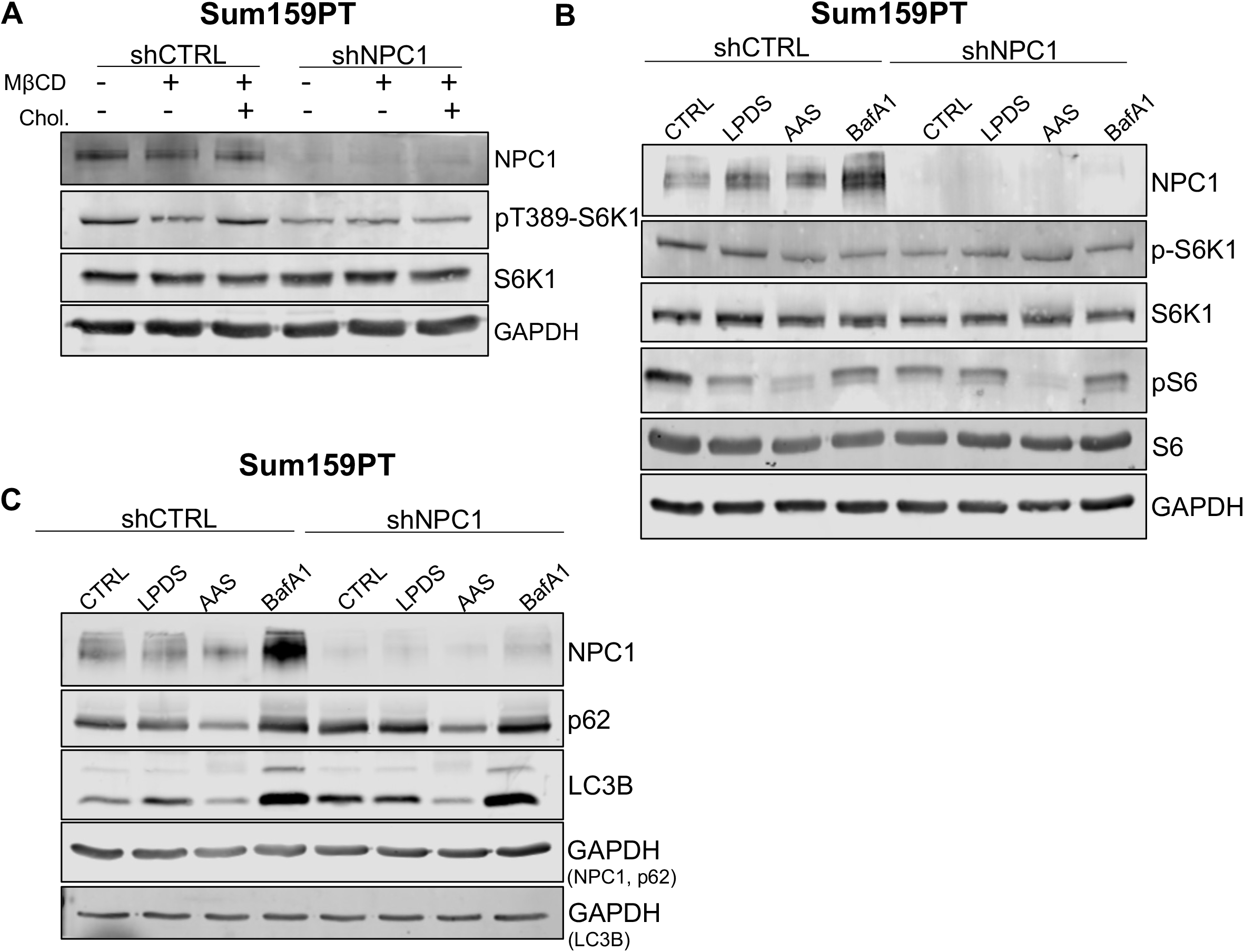
NPC1 affects mTOR and autophagy signaling under stress conditions in TNBC. **A)** Western of phospho-S6 kinase signaling under cholesterol-stressed conditions. Control cells (lane 1,4) are maintained in full serum. Where indicated, MβCD is used to deplete cholesterol for 2hr and then, as indicated (lane 3, 6), MβCD is removed and replaced with MβCD:Cholesterol to replenish cholesterol supply. **B)** S6 kinase signaling under metabolic or lysosome stress conditions for 24 hours (LPDS-5% Lipoprotein depleted serum; AAS-amino acid depletion with 5% FBS; BafA1-5nM). **C)** Autophagy markers p62 and LC3B under metabolic or lysosomal stress conditions.

To further evaluate the effect of NPC1 silencing on mTOR, Sum159PT cells were treated in metabolic or lysosomal-stress conditions for 24 hours. Knockdown of NPC1 caused a reduction in p-S6K1 and p-S6 in full serum and nutrient conditions (**Figure 5B**). Under nutrient and lysosomal stress, S6K signaling remained roughly the same between control and knockdown cells. However, nutrient deprivation coupled with NPC1 silencing caused further reduction of 4EBP-1 signaling than nutrient deprivation alone (**Supplementary 7B**). Together, these data suggest NPC1 silencing affects S6K versus 4EBP1 nodes of mTOR differently depending on nutrient status.

The effects of NPC1 on lysosomal metabolism and mTOR signaling implicates autophagy in this model, and autophagy defects have been noted in Niemann-Pick disease models. Indeed, NPC1 silencing causes increased LC3-II at baseline and in response to metabolic or lysosomal stress (**Figure 5C**), suggesting either increased production of autophagosomes or the buildup of autophagosomes. p62, which accumulates when autophagic clearance is impaired, was also increased in NPC1 knockdown cells. Together, these data are consistent with Niemann-Pick models where autophagosomes can fuse with the lysosomes but cannot be properly degraded.

Together, these data demonstrate that NPC1 silencing acts on the mTOR pathway in association with decreased cancer cell viability, proliferation, and invasive capacity. Further, mitochondrial defects accompany this overall phenotype, and cells become less able to survive clinically relevant anti-cancer drugs.

### NPC1 has therapeutic potential alone and in combination with chemotherapy in TNBC

We have shown that NPC1 silencing in TNBC causes mitochondrial dysfunction, decreased proliferation and invasion, and suppression of mTOR signaling (**Figure 6A**). Together this suggests NPC1 as a potential target in this cancer subtype. Specific NPC1 inhibitors are not commercially available, although several small-molecule compounds have been shown to directly bind and inhibit NPC1 in addition to other targets with sterol-sensing domains^(42,43)^, most notably U18666A. To evaluate U18666A as a potential therapeutic, three TNBC cell lines were treated with serial dilutions of drug for 48 hours, which caused reduced cell viability at micromolar doses (**Figure 6B**).

**Figure 6:**
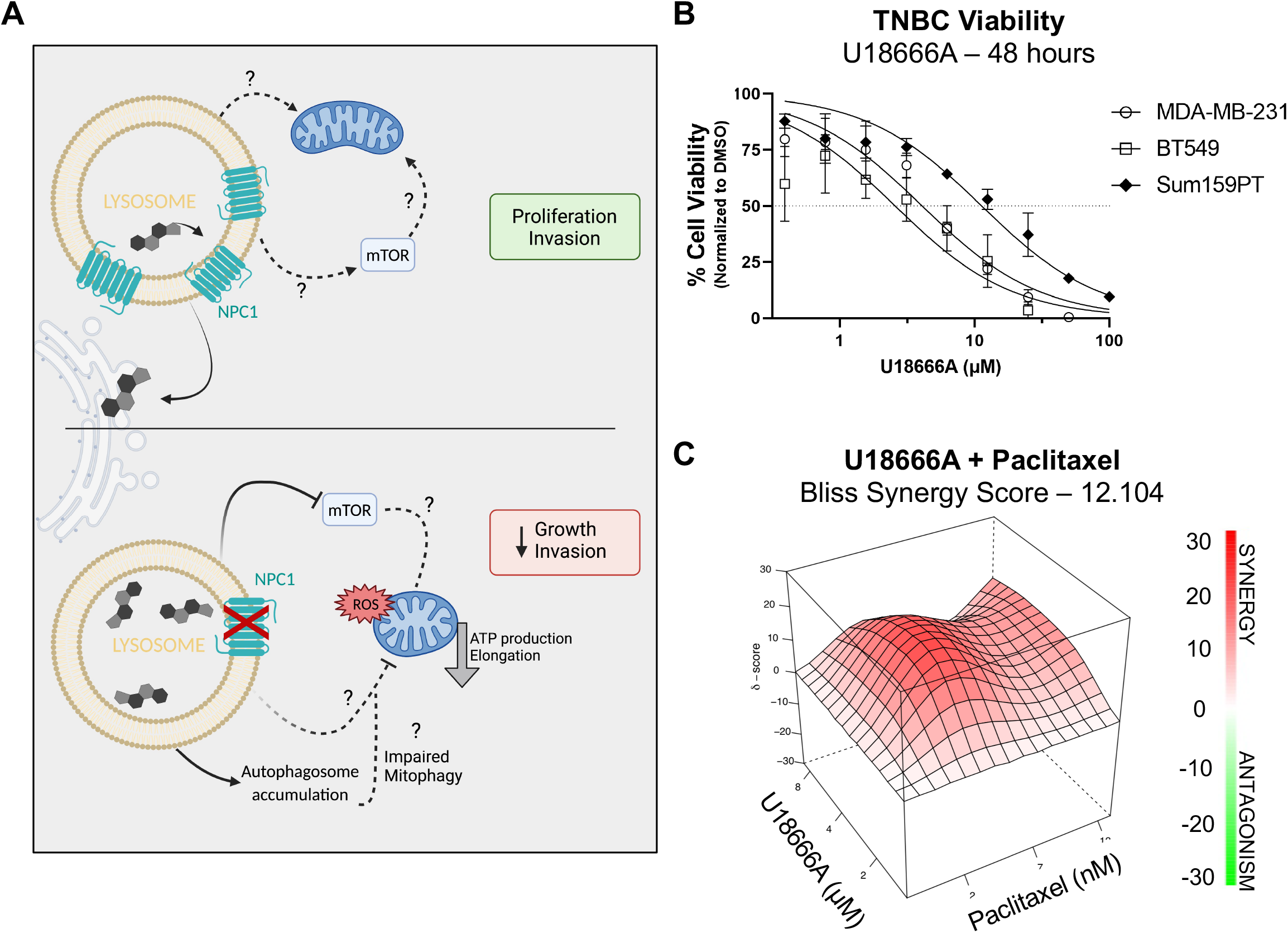
NPC1 has therapeutic potential alone and in combination with chemotherapies in TNBC. **A)** Graphical summary of the anti-tumor effects of NPC1 silencing. Created with biorender.com **B)** Dose response curves of U18666A in three TNBC cell lines, 48 hours drug treatment as analyzed by crystal violet. Linear regression with variable slope. **C)** Additive and synergistic effects of U18666A and Paclitaxel in BT549 cells. 3 doses of each drug were given for 48 hours and analyzed by crystal violet. Additive/Synergy effect analyzed and calculated using SynergyFinder 2.0.

Because chemotherapy remains the frontline therapeutic in TNBC, U18666A was combined with paclitaxel. This led to additive or synergistic effect^(44)^, depending on dose combinations (**Figure 6C, Supplementary 7C**). These data demonstrate the potential utility of targeting NPC1 alone or in combination with existing TNBC therapies.

## DISCUSSION

TNBC is an aggressive breast cancer subtype with limited therapeutic options. The EMT-suppressor miR-200c is lost due to silencing or deletion in TNBC, allowing a phenotypic switch that supports invasion and metastasis^(45)^. Using miR-200c as a tool to identify novel pathways upregulated in TNBC, we found that miR-200c directly regulates *NPC1*, a lysosomal cholesterol transporter that we find to be elevated in TNBC compared to ER positive disease. Literature evaluating NPC1 in cancer is extremely limited, with few studies showing elevated NPC1 in certain cancers, and implicating NPC1 in cell invasion and cell growth^(24,46,47)^. In our study, knockdown of NPC1 in three TNBC cell lines led to significant loss of proliferation in 2D culture and colony formation in soft agar. The complexity of NPC1 function in normal physiology suggests numerous mechanisms by which NPC1 could provide a survival advantage for TNBC, including support of cell proliferation and viability, cholesterol homeostasis, lysosomal function, and known interactions with mTOR signaling.

Because NPC1 is a major provider of exogenous cholesterol to the endoplasmic reticulum, we hypothesized that disrupting this cholesterol axis could prevent cells from obtaining adequate exogenous cholesterol. Decreased growth in LPDS conditions existed within a subset of breast cancer cell lines and occurred more frequently in TNBC cell lines as compared to those representing other breast cancer subtypes. Demand for exogenous cholesterol weakly but significantly correlates with NPC1 protein expression, but not mRNA levels of other cholesterol-related genes. This suggests that while cholesterol auxotrophy exists within breast cancer, it may not be driven by a specific mechanism that is shared among all breast cancer auxotrophs.

It is possible that cholesterol auxotrophic breast cancers have elevated NPC1 protein as a mechanism to support exogenous cholesterol availability. However, cell-permeable cholesterol did not rescue the viability of NPC1 knockdown TNBC cells in soft-agar conditions, suggesting that the slowed growth in these cells is not primarily driven by loss of lysosomal-endoplasmic reticulum cholesterol transport. Further, TNBC cells did not exhibit an induction of sterol-related genes that generally respond to low-cholesterol levels, suggesting cells received adequate intracellular cholesterol without the contribution of NPC1 in 2D and 3D culture. This is likely due to the existence of unique transporters that transfer cholesterol to other organelles, such as the lysosome-mitochondrial transporter STARD3 which is implicated in Niemann-Pick disease^(48)^. Therefore, while NPC1 loss causes accumulation of cholesterol within the lysosomes, which is confirmed by immunocytochemistry of cholesterol/filipin staining, it does not completely block access of all organelles to lysosomal cholesterol pools.

Having ruled out cholesterol depletion as a primary mediator of the effects of NPC1 silencing growth inhibition, we evaluated mitochondrial metabolism. NPC1 depletion led to a loss of mitochondrial respiration and increase in mitochondrial ROS, which was associated with shortened and rounded mitochondria. Morphology of mitochondria is largely dictated by fission and fusion processes^(49)^. In the context of cancer, increased fission is believed to aid in invasion and migration, as shortened mitochondria can be more easily trafficked to the leading edge and to areas of high energy demand^(50,51)^. This is inconsistent with the finding that NPC1 knockdown cells had decreased invasive capacity along with shortened mitochondria. However, we observed numerous “donut” mitochondria in NPC1 cells, which are believed to be caused by abnormal mitochondrial calcium signaling or ROS levels and have been noted to be less efficient in ATP production^(52)^. This suggests that the shortening of mitochondria in these cells may be driven by mitochondrial stress, and that these shorter defective mitochondria are not able to support the high energetic demands of tumor cell invasion. A limitation of our studies is that we do not know the precise mechanisms driving altered mitochondrial morphology and function downstream of NPC1 in TNBC, which warrants future investigation. Interestingly, mitochondrial cholesterol content is increased in some Niemann-Pick models^(53,54)^, while in others, mitochondrial biogenesis is suppressed^(36)^ or mitophagy is defective^(35)^. Our novel finding that NPC1 silencing can suppress P70-S6K signaling in some breast cancer cell lines suggests that mTOR may link lysosomal cholesterol with mitochondria in these cells.

Together, these data characterize for the first time the effect of NPC1 inhibition in TNBC, and propose a role for this protein in cancer. NPC1 expression is elevated in TNBC, and genetic inhibition of NPC1 caused decreased cell proliferation, growth on soft agar, and decreased invasive capacity. This was associated with observed mitochondrial defects, and suppressed mTOR signaling. We also find that U18666A, a small molecule that inhibits NPC1 activity, has low micromolar IC50 against TNBC cell lines, as well as synergy in combination with the clinically approved chemotherapeutic, paclitaxel. Thus, targeting NPC1 could be of therapeutic interest alone or in combination with standard treatments in TNBC.

## Supporting information

Supplementary - primer sequences

## ACKNOWLEDGEMENTS

We acknowledge the University of Colorado Cancer Center Support Grant P30CA046934, particularly the Functional Genomics and Cell Technologies Shared Recourses. Seahorse assays were done in partnership with the CU Anschutz NORC Core (P30 DK048520/DK/NIDDK NIH HHS/United States). Microscopy was performed at the Advanced Light Microscopy Core-NeuroTechnology Center at University of Colorado Anschutz Medical Campus, supported in part by Rocky Mountain Neurological Disorders Core Grant Number P30 NS048154 and by Diabetes Research Center Grant Number P30 DK116073. The authors thank Dr. Roberto Zoncu at University of California, Berkeley for NPC1 constructs and Dr. Christina Towers at Salk Institute for autophagy guidance. The graphical abstract in figure 6 was created using Biorender under agreement #MB23UM4TEF.

## REFERENCES

1. Abramson VG, Lehmann BD, Ballinger TJ, Pietenpol JA. Subtyping of triple-negative breast cancer: implications for therapy. Cancer 2015;121(1):8–16 doi 10.1002/cncr.28914.

2. Liedtke C, Mazouni C, Hess KR, André F, Tordai A, Mejia JA, et al. Response to Neoadjuvant Therapy and Long-Term Survival in Patients With Triple-Negative Breast Cancer. Journal of Clinical Oncology 2008;26(8):1275–81 doi 10.1200/JCO.2007.14.4147.

3. Cochrane DR, Cittelly DM, Howe EN, Spoelstra NS, McKinsey EL, LaPara K, et al. MicroRNAs Link Estrogen Receptor Alpha Status and Dicer Levels in Breast Cancer. Hormones and Cancer 2010;1(6):306–19 doi 10.1007/s12672-010-0043-5.

4. Feng Z-M, Qiu J, Chen X-W, Liao R-X, Liao X-Y, Zhang L-P, et al. Essential role of miR-200c in regulating self-renewal of breast cancer stem cells and their counterparts of mammary epithelium. BMC Cancer 2015;15(1):645 doi 10.1186/s12885-015-1655-5.

5. Lim Y-Y, Wright JA, Attema JL, Gregory PA, Bert AG, Smith E, et al. Epigenetic modulation of the miR-200 family is associated with transition to a breast cancer stem-cell-like state. Journal of Cell Science 2013;126(10):2256–66 doi 10.1242/jcs.122275.

6. Howe EN, Cochrane DR, Richer JK. Targets of miR-200c mediate suppression of cell motility and anoikis resistance. Breast cancer research : BCR 2011;13(2):R45–R doi 10.1186/bcr2867.

7. Burk U, Schubert J, Wellner U, Schmalhofer O, Vincan E, Spaderna S, et al. A reciprocal repression between ZEB1 and members of the miR-200 family promotes EMT and invasion in cancer cells. EMBO reports 2008;9(6):582–9 doi 10.1038/embor.2008.74.

8. Kawaguchi T, Yan L, Qi Q, Peng X, Gabriel EM, Young J, et al. Overexpression of suppressive microRNAs, miR-30a and miR-200c are associated with improved survival of breast cancer patients. Scientific Reports 2017;7(1):15945 doi 10.1038/s41598-017-16112-y.

9. Kalluri R, Weinberg RA. The basics of epithelial-mesenchymal transition. The Journal of Clinical Investigation 2009;119(6):1420–8 doi 10.1172/JCI39104.

10. Mak MP, Tong P, Diao L, Cardnell RJ, Gibbons DL, William WN, et al. A Patient-Derived, Pan-Cancer EMT Signature Identifies Global Molecular Alterations and Immune Target Enrichment Following Epithelial-to-Mesenchymal Transition. Clinical Cancer Research 2016;22(3):609–20 doi 10.1158/1078-0432.CCR-15-0876.

11. Polytarchou C, Iliopoulos D, Struhl K. An integrated transcriptional regulatory circuit that reinforces the breast cancer stem cell state. Proceedings of the National Academy of Sciences of the United States of America 2012;109(36):14470–5 doi 10.1073/pnas.1212811109.

12. Rogers TJ, Christenson JL, Greene LI, Neill KI, Williams MM, Gordon MA, et al. Reversal of Triple-negative Breast Cancer EMT by miR-200c Decreases Tryptophan Catabolism and a Program of Immune-Suppression. Molecular Cancer Research 2018.

13. Williams MM, Hafeez SA, Christenson JL, O’Neill KI, Hammond NG, Richer JK. Reversing an Oncogenic Epithelial-to-Mesenchymal Transition Program in Breast Cancer Reveals Actionable Immune Suppressive Pathways. Pharmaceuticals (Basel, Switzerland) 2021;14(11) doi 10.3390/ph14111122.

14. Goldstein JL, Brown MS. A century of cholesterol and coronaries: from plaques to genes to statins. Cell 2015;161(1):161–72 doi 10.1016/j.cell.2015.01.036.

15. Li X, Wang J, Coutavas E, Shi H, Hao Q, Blobel G. Structure of human Niemann–Pick C1 protein. Proceedings of the National Academy of Sciences 2016;113(29):8212–7 doi 10.1073/pnas.1607795113.

16. Kennedy BE, Madreiter CT, Vishnu N, Malli R, Graier WF, Karten B. Adaptations of energy metabolism associated with increased levels of mitochondrial cholesterol in Niemann-Pick type C1-deficient cells. The Journal of biological chemistry 2014;289(23):16278–89 doi 10.1074/jbc.M114.559914.

17. Geberhiwot T, Moro A, Dardis A, Ramaswami U, Sirrs S, Marfa MP, et al. Consensus clinical management guidelines for Niemann-Pick disease type C. Orphanet Journal of Rare Diseases 2018;13(1):50 doi 10.1186/s13023-018-0785-7.

18. Castellano BM, Thelen AM, Moldavski O, Feltes M, van der Welle REN, Mydock-McGrane L, et al. Lysosomal cholesterol activates mTORC1 via an SLC38A9–Niemann-Pick C1 signaling complex. Science 2017;355(6331):1306–11 doi 10.1126/science.aag1417.

19. Lim C-Y, Davis OB, Shin HR, Zhang J, Berdan CA, Jiang X, et al. ER–lysosome contacts enable cholesterol sensing by mTORC1 and drive aberrant growth signalling in Niemann–Pick type C. Nature Cell Biology 2019;21(10):1206–18 doi 10.1038/s41556-019-0391-5.

20. Gévry N, Schoonjans K, Guay F, Murphy BD. Cholesterol supply and SREBPs modulate transcription of the Niemann-Pick C-1 gene in steroidogenic tissues. Journal of Lipid Research 2008;49(5):1024–33 doi https://doi.org/10.1194/jlr.M700554-JLR200.

21. Garver WS, Jelinek D, Francis GA, Murphy BD. The Niemann-Pick C1 gene is downregulated by feedback inhibition of the SREBP pathway in human fibroblasts. Journal of lipid research 2008;49(5):1090–102 doi 10.1194/jlr.M700555-JLR200.

22. Gévry NY, Lalli E, Sassone-Corsi P, Murphy BD. Regulation of Niemann-Pick C1 Gene Expression by the 3′5′-Cyclic Adenosine Monophosphate Pathway in Steroidogenic Cells. Molecular Endocrinology 2003;17(4):704–15 doi 10.1210/me.2002-0093.

23. Nam JW, Rissland OS, Koppstein D, Abreu-Goodger C, Jan CH, Agarwal V, et al. Global analyses of the effect of different cellular contexts on microRNA targeting. Mol Cell 2014;53(6):1031–43 doi 10.1016/j.molcel.2014.02.013.

24. Jose J, Hoque M, Engel J, Beevi SS, Wahba M, Georgieva MI, et al. Annexin A6 and NPC1 regulate LDL-inducible cell migration and distribution of focal adhesions. Scientific Reports 2022;12(1):596 doi 10.1038/s41598-021-04584-y.

25. Das A, Brown MS, Anderson DD, Goldstein JL, Radhakrishnan A. Three pools of plasma membrane cholesterol and their relation to cholesterol homeostasis. eLife 2014;3 doi 10.7554/eLife.02882.

26. Nieweg K, Schaller H, Pfrieger FW. Marked differences in cholesterol synthesis between neurons and glial cells from postnatal rats. Journal of Neurochemistry 2009;109(1):125–34 doi 10.1111/j.1471-4159.2009.05917.x.

27. Garcia-Bermudez J, Baudrier L, Bayraktar EC, Shen Y, La K, Guarecuco R, et al. Squalene accumulation in cholesterol auxotrophic lymphomas prevents oxidative cell death. Nature 2019;567(7746):118–22 doi 10.1038/s41586-019-0945-5.

28. Riscal R, Bull CJ, Mesaros C, Finan JM, Carens M, Ho ES, et al. Cholesterol Auxotrophy as a Targetable Vulnerability in Clear Cell Renal Cell Carcinoma. Cancer discovery 2021;11(12):3106 doi 10.1158/2159-8290.CD-21-0211.

29. Abi-Mosleh L, Infante Rodney E, Radhakrishnan A, Goldstein Joseph L, Brown Michael S. Cyclodextrin overcomes deficient lysosome-to-endoplasmic reticulum transport of cholesterol in Niemann-Pick type C cells. Proceedings of the National Academy of Sciences 2009;106(46):19316–21 doi 10.1073/pnas.0910916106.

30. Brogden G, Shammas H, Walters F, Maalouf K, Das AM, Naim HY, et al. Different Trafficking Phenotypes of Niemann-Pick C1 Gene Mutations Correlate with Various Alterations in Lipid Storage, Membrane Composition and Miglustat Amenability. Int J Mol Sci 2020;21(6) doi 10.3390/ijms21062101.

31. Frolov A, Zielinski SE, Crowley JR, Dudley-Rucker N, Schaffer JE, Ory DS. NPC1 and NPC2 Regulate Cellular Cholesterol Homeostasis through Generation of Low Density Lipoprotein Cholesterol-derived Oxysterols*. Journal of Biological Chemistry 2003;278(28):25517–25 doi https://doi.org/10.1074/jbc.M302588200.

32. DeBose-Boyd RA, Ye J. SREBPs in Lipid Metabolism, Insulin Signaling, and Beyond. Trends in biochemical sciences 2018;43(5):358–68 doi 10.1016/j.tibs.2018.01.005.

33. Wei J, Zhang YY, Luo J, Wang JQ, Zhou YX, Miao HH, et al. The GARP Complex Is Involved in Intracellular Cholesterol Transport via Targeting NPC2 to Lysosomes. Cell Rep 2017;19(13):2823–35 doi 10.1016/j.celrep.2017.06.012.

34. Naito T, Ercan B, Krshnan L, Triebl A, Koh DHZ, Wei F-Y, et al. Movement of accessible plasma membrane cholesterol by the GRAMD1 lipid transfer protein complex. eLife 2019;8:e51401 doi 10.7554/eLife.51401.

35. Davis OB, Shin HR, Lim CY, Wu EY, Kukurugya M, Maher CF, et al. NPC1-mTORC1 Signaling Couples Cholesterol Sensing to Organelle Homeostasis and Is a Targetable Pathway in Niemann-Pick Type C. Dev Cell 2021;56(3):260-76.e7 doi 10.1016/j.devcel.2020.11.016.

36. Yambire KF, Fernandez-Mosquera L, Steinfeld R, Mühle C, Ikonen E, Milosevic I, et al. Mitochondrial biogenesis is transcriptionally repressed in lysosomal lipid storage diseases. eLife 2019;8:e39598 doi 10.7554/eLife.39598.

37. Sundqvist M, Christenson K, Björnsdottir H, Osla V, Karlsson A, Dahlgren C, et al. Elevated Mitochondrial Reactive Oxygen Species and Cellular Redox Imbalance in Human NADPH-Oxidase-Deficient Phagocytes. Frontiers in Immunology 2017;8 doi 10.3389/fimmu.2017.01828.

38. Miyazono Y, Hirashima S, Ishihara N, Kusukawa J, Nakamura K-i, Ohta K. Uncoupled mitochondria quickly shorten along their long axis to form indented spheroids, instead of rings, in a fission-independent manner. Scientific Reports 2018;8(1):350 doi 10.1038/s41598-017-18582-6.

39. Picard M, McEwen Bruce S. Mitochondria impact brain function and cognition. Proceedings of the National Academy of Sciences 2014;111(1):7–8 doi 10.1073/pnas.1321881111.

40. Caino MC, Altieri DC. Molecular Pathways: Mitochondrial Reprogramming in Tumor Progression and Therapy. Clinical cancer research : an official journal of the American Association for Cancer Research 2016;22(3):540–5 doi 10.1158/1078-0432.Ccr-15-0460.

41. Shah SP, Roth A, Goya R, Oloumi A, Ha G, Zhao Y, et al. The clonal and mutational evolution spectrum of primary triple-negative breast cancers. Nature 2012;486(7403):395–9 doi 10.1038/nature10933.

42. Lu F, Liang Q, Abi-Mosleh L, Das A, De Brabander JK, Goldstein JL, et al. Identification of NPC1 as the target of U18666A, an inhibitor of lysosomal cholesterol export and Ebola infection. eLife 2015;4:e12177 doi 10.7554/eLife.12177.

43. Quan X, Chen X, Sun D, Xu B, Zhao L, Shi X, et al. The mechanism of the effect of U18666a on blocking the activity of 3β-hydroxysterol Δ-24-reductase (DHCR24): molecular dynamics simulation study and free energy analysis. Journal of Molecular Modeling 2016;22(2):46 doi 10.1007/s00894-016-2907-2.

44. Ianevski A, Giri AK, Aittokallio T. SynergyFinder 2.0: visual analytics of multi-drug combination synergies. Nucleic Acids Research 2020;48(W1):W488-W93 doi 10.1093/nar/gkaa216.

45. Dongre A, Weinberg RA. New insights into the mechanisms of epithelial–mesenchymal transition and implications for cancer. Nature Reviews Molecular Cell Biology 2019;20(2):69–84 doi 10.1038/s41580-018-0080-4.

46. Head SA, Shi WQ, Yang EJ, Nacev BA, Hong SY, Pasunooti KK, et al. Simultaneous Targeting of NPC1 and VDAC1 by Itraconazole Leads to Synergistic Inhibition of mTOR Signaling and Angiogenesis. ACS chemical biology 2017;12(1):174–82 doi 10.1021/acschembio.6b00849.

47. Lyu J, Yang EJ, Head SA, Ai N, Zhang B, Wu C, et al. Pharmacological blockade of cholesterol trafficking by cepharanthine in endothelial cells suppresses angiogenesis and tumor growth. Cancer Lett 2017;409:91–103 doi 10.1016/j.canlet.2017.09.009.

48. Höglinger D, Burgoyne T, Sanchez-Heras E, Hartwig P, Colaco A, Newton J, et al. NPC1 regulates ER contacts with endocytic organelles to mediate cholesterol egress. Nature Communications 2019;10(1):4276 doi 10.1038/s41467-019-12152-2.

49. Westermann B. Mitochondrial fusion and fission in cell life and death. Nature Reviews Molecular Cell Biology 2010;11(12):872–84 doi 10.1038/nrm3013.

50. Boulton DP, Caino MC. Mitochondrial Fission and Fusion in Tumor Progression to Metastasis. Front Cell Dev Biol 2022;10 doi 10.3389/fcell.2022.849962.

51. Trotta AP, Chipuk JE. Mitochondrial dynamics as regulators of cancer biology. Cellular and Molecular Life Sciences 2017;74(11):1999–2017 doi 10.1007/s00018-016-2451-3.

52. Liu X, Hajnóczky G. Altered fusion dynamics underlie unique morphological changes in mitochondria during hypoxia–reoxygenation stress. Cell Death & Differentiation 2011;18(10):1561–72 doi 10.1038/cdd.2011.13.

53. Yu W, Gong J-S, Ko M, Garver WS, Yanagisawa K, Michikawa M. Altered Cholesterol Metabolism in Niemann-Pick Type C1 Mouse Brains Affects Mitochondrial Function *<sup></sup>. Journal of Biological Chemistry 2005;280(12):11731–9 doi 10.1074/jbc.M412898200.

54. Charman M, Kennedy BE, Osborne N, Karten B. MLN64 mediates egress of cholesterol from endosomes to mitochondria in the absence of functional Niemann-Pick Type C1 protein. Journal of Lipid Research 2010;51(5):1023–34 doi https://doi.org/10.1194/jlr.M002345.

